# An MR-based brain template and atlas for optical projection tomography and light sheet fluorescence microscopy

**DOI:** 10.1101/2022.11.14.516420

**Authors:** Stefanie M.A. Willekens, Federico Morini, Tomas Mediavilla, Emma Nilsson, Greger Orädd, Max Hahn, Nunya Chotiwan, Montse Visa, Per-Olof Berggren, Erwin Ilegemns, Anna K. Överby, Ulf Ahlgren, Daniel Marcellino

## Abstract

Optical projection tomography (OPT) and light sheet fluorescence microscopy (LSFM) are high-resolution optical imaging techniques operating in the mm-cm range, ideally suited for *ex vivo* 3D whole mouse brain imaging. Although these techniques exhibit high sensitivity and specificity for antibody-labeled targets, the provided anatomical information remains limited. To allow anatomical mapping of fluorescent signal in whole brain, we developed a novel magnetic resonance (MR) – based template with its associated tissue priors and atlas labels, specifically designed for brains subjected to tissue processing protocols required for 3D optical imaging. We investigated the effect of tissue pre-processing and clearing on brain size and morphology and developed optimized templates for BABB/Murrays clear (OCUM) and DBE/iDISCO (iOCUM) cleared brains. By creating optical-(i)OCUM fusion images using our mapping procedure, we localized dopamine transporter and translocator protein expression and tracer innervation from the eye to the lateral geniculate nucleus of thalamus and superior colliculus. These fusion images allowed for precise anatomical identification of fluorescent signal in discrete brain areas. As such, these templates enable applications in a broad range of research areas integrating optical 3D brain imaging by providing an MR template for cleared brains.

## Introduction

Three-dimensional (3D) visualization of specific cell populations, protein expression patterns or pathologic markers at the level of the whole brain represents an invaluable tool in neuroscience. Optical projection tomography (OPT) and light sheet fluorescence microscopy (LSFM) are high-resolution optical 3D imaging techniques, enabling the visualization of specifically labeled targets in mesoscopic sized (mm-cm range) transparent specimens^1,2^. Therefore, these optical techniques harbor great suitability for *ex vivo* whole rodent brain imaging, providing information at cellular resolution in the intact brain^3,4^. In line with other functional imaging modalities, OPT and LSFM display high sensitivity and specificity for their target, but offer only limited anatomical information. Considering the highly compartmentalized anatomy of the brain^5^ and the specific roles these regions fulfill, it is of the utmost importance to be able to map the fluorescent signals, acquired by OPT or LSFM, to annotated brain regions. The possibility to anatomically map protein expression profiles and perform 3D quantification and statistics on these images, would greatly benefit the application of optical mesoscopic imaging in neuroscience. The first step towards these analyses is to co-register the optical brain signals to a reference brain for which detailed annotated brain regions can be readily identified.

The Common Coordinate Framework version 3 (CCFv3) of the Allen Institute of Brain Sciences (AIBS)^5-7^ has been used for co-registration and quantitative analyses of 3D optical brain images^8,9^ and has formed the basis for LSFM-specific brain templates based on optically cleared brains, which were successfully applied to study drug effects in the whole brain^10,11^. Since the AIBS template and concordant atlas are based on serial two-photon tomography (STPT) imaging^5^ morphometric discrepancies can be observed mainly in the most rostral and caudal regions, when compared to whole brain MRI. Nevertheless, its applicability has been demonstrated well suited to link connectivity patterns, functional properties and cellular architecture^6,12^ The LSFM-based templates, on the other hand, are based on tissue autofluorescence and therefore may generate suboptimal tissue contrast and anatomical detail to distinguish all brain regions. Magnetic Resonance Imaging (MRI) is well known to generate detailed anatomical brain images due to its high resolution and exquisite tissue contrast. Therefore, MR images are ideally suited as an anatomical reference for the creation of fusion images by means of co-registration. The most straightforward way to create fluorescence-MR fusion images is by using an MRI-based mouse brain template, of which several are currently available^13-15^. Whereas the AIBS template is based on histology and the forementioned MRI-based templates originate from MR scans acquired either *in vivo* or *ex vivo in situ* (acquired within the skull), OPT and LSFM images are acquired after brain removal from the skull and following extensive processing and tissue clearing. These processes are known to exert differential effects on brain size and morphology dependent on which tissue clearing protocol is applied^16^.

Here, we present the creation of a novel high-resolution (40 μm^3^ isotropic voxel size) Optically Cleared UMeå (OCUM) brain template, consisting of *ex vivo* T1-weighted MR images acquired after tissue processing and clearing for optical imaging, with its associated tissue priors and corresponding atlas annotating 336 regions of interest (ROIs). Furthermore, two distinct versions of the template and its corresponding atlas are presented, each optimized for two distinct clearing methods that have differential effects on brain size after clearing and rehydration for MRI acquisition. Finally, we demonstrate the utility of OCUM by creating fluorescence-template fusion images with 3D optical images visualizing the dopamine transporter (DAT), the 18 kDa translocator protein (TSPO), and optic nerve innervation of the lateral geniculate nucleus (LGN) of thalamus and superior colliculus. In each case, the fusion images allowed for precise anatomical identification of brain regions with fluorescent signal. As such, these templates will significantly enhance the applicability of mesoscopic optical imaging in neuroscience.

## Results

### Inadequate anatomical referencing with autofluorescence

It was recently shown that significant information can be obtained from tissue autofluorescence^17^ using OPT/LSFM and, it is well known that the brain generally displays a high level of autofluorescence. Therefore, we first aimed to optimize the reconstruction of the autofluorescence signal, acquired from OPT, to obtain more anatomical details and co-registered the dopamine transporter signal to the improved autofluorescence signal (figure 1). Although we were able to obtain improved tissue contrast between the grey matter (GM) and white matter (WM) regions and improved delineation of the deep nuclei, anatomical detail remained insufficient to identify detailed anatomical regions, mainly in cortical areas. In theory, this contrast could be further improved by enhancing and optimizing the autofluorescence signal. However, to obtain high signal to noise ratio’s for the antibody-target optical signals, autofluorescence should remain as limited as possible. Therefore, accurate anatomical mapping of 3D optical brain signals cannot be obtained from tissue autofluorescence.

**Figure 1:**
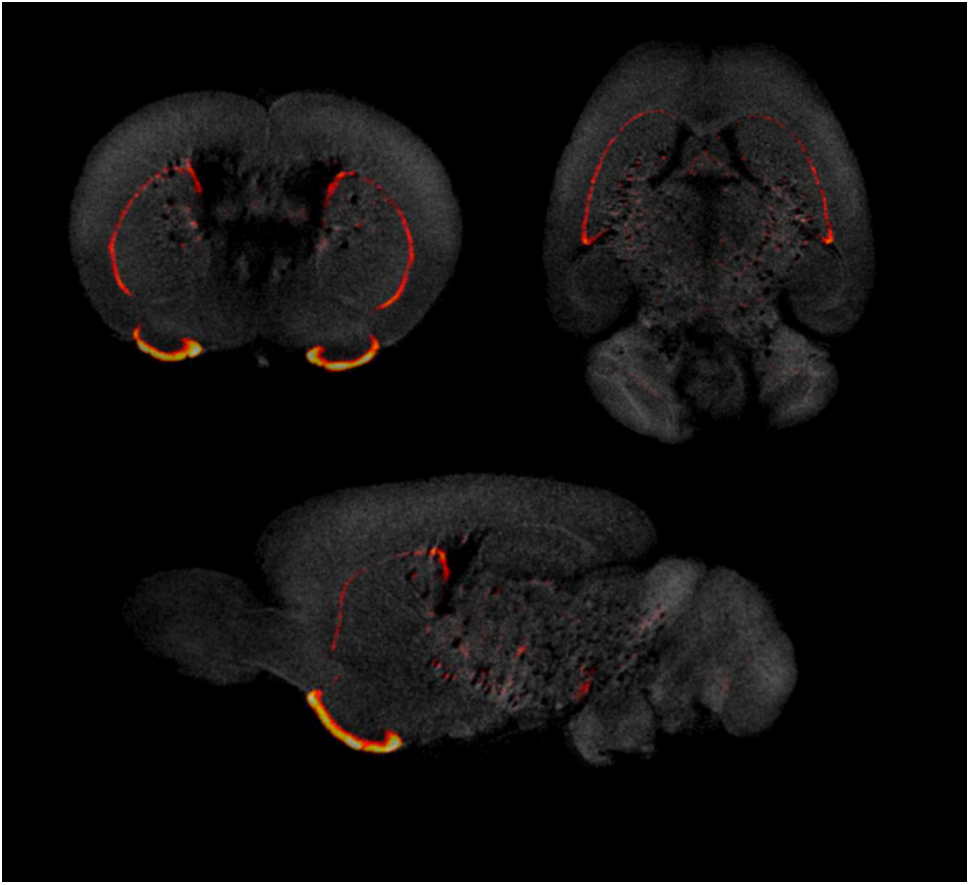
Insufficient anatomical mapping based on brain autofluorescence. Co-registration of DAT signal acquired by OPT with the anatomy, reconstructed based on the tissues autofluorescence, provides insufficient anatomical detail for brain.

### Differential effects of clearing agents on brain volume

Apart from the anatomical reference problem exemplified in figure 1, brain clearing protocols are known to exert extensive effects on brain morphology which can be differential in distinct brain regions^16^. To obtain a representative anatomical reference for optically cleared brains with these morphological changes, we acquired *ex vivo* structural T1-weighted images of brains that were subjected to tissue processing and optical clearing with benzyl alcohol-benzyl benzoate (BABB)/Murray’s clear (n=10)^1,18^ or with dibenzyl ether (DBE)/iDISCO (n=9)^19^, two frequently used protocols for optical tissue clearing. To investigate the effects of both BABB and DBE clearing on brain size and morphology, we initially created two individual brain templates, namely one for BABB cleared brains and one for DBE cleared brains, respectively. The individual templates were calculated as the mid-point average from the serial registration of all individual, bias-corrected MR-images. The head-to-head comparison of the average BABB and DBE templates revealed a clear difference in size, for which the BABB template was significantly larger than the DBE template (Figure 2a and b). Furthermore, the size difference could not be attributed to sole differences in cortical shrinkage since a clear mismatch was observed in WM tracts and deep nuclei when superimposed in an identical image space (Figure 2a and b). Segment-based brain volume calculations on the individual T1-weighted images revealed that DBE-cleared brains (0.308 cm^3^ ± 0.009) were significantly smaller when compared to BABB-cleared brains (0.483 ± 0.023 cm^3^) (Figure 2c). To further characterize the effect of tissue processing and clearing methods on the brain, we proceeded by comparing these volumes with brain volumes calculated from T1-weighted scans acquired *in vivo* (n=62)^20^ and *ex vivo in situ* (n=40), as well as with brain volumes based on tissue autofluorescence of individual BABB and DBE brains (Figure 2c). We observed significant differences between *in vivo* brain volume (0.461 ± 0.011 cm^3^) and all other tested volumes. On T1-weighted images, both the *ex vivo (*0.437 ± 0.011 cm^3^) and DBE brains (0.308 ± 0.01 cm^3^) were significantly smaller (p<0.001) than the *in vivo* brain volume, while the BABB brain volume was significantly larger (0.483 ± 0.023 cm^3^; p=0.03); Figure 2c). However, when BABB (0.343 ± 0.0517 cm^3^) and DBE (0.299 ± 0.071 cm^3^) brain volumes were calculated from tissue autofluorescence, they were not significantly different (p=0.12). Interestingly, BABB brain volume was significantly larger when calculated based on T1-weighted images (0.483 ± 0.023 cm^3^) as when calculated from tissue autofluorescence (0.343 ± 0.0517 cm^3^; p<0.001), which was not the case for the DBE cleared brains (p=0.94), indicating a distinct effect of BABB and DBE clearing on brain rehydration, respectively (Figure 2c). Together, these particular volume effects due to dehydration, rehydration and clearing plead for the application of a representative, optically cleared brain template.

**Figure 2:**
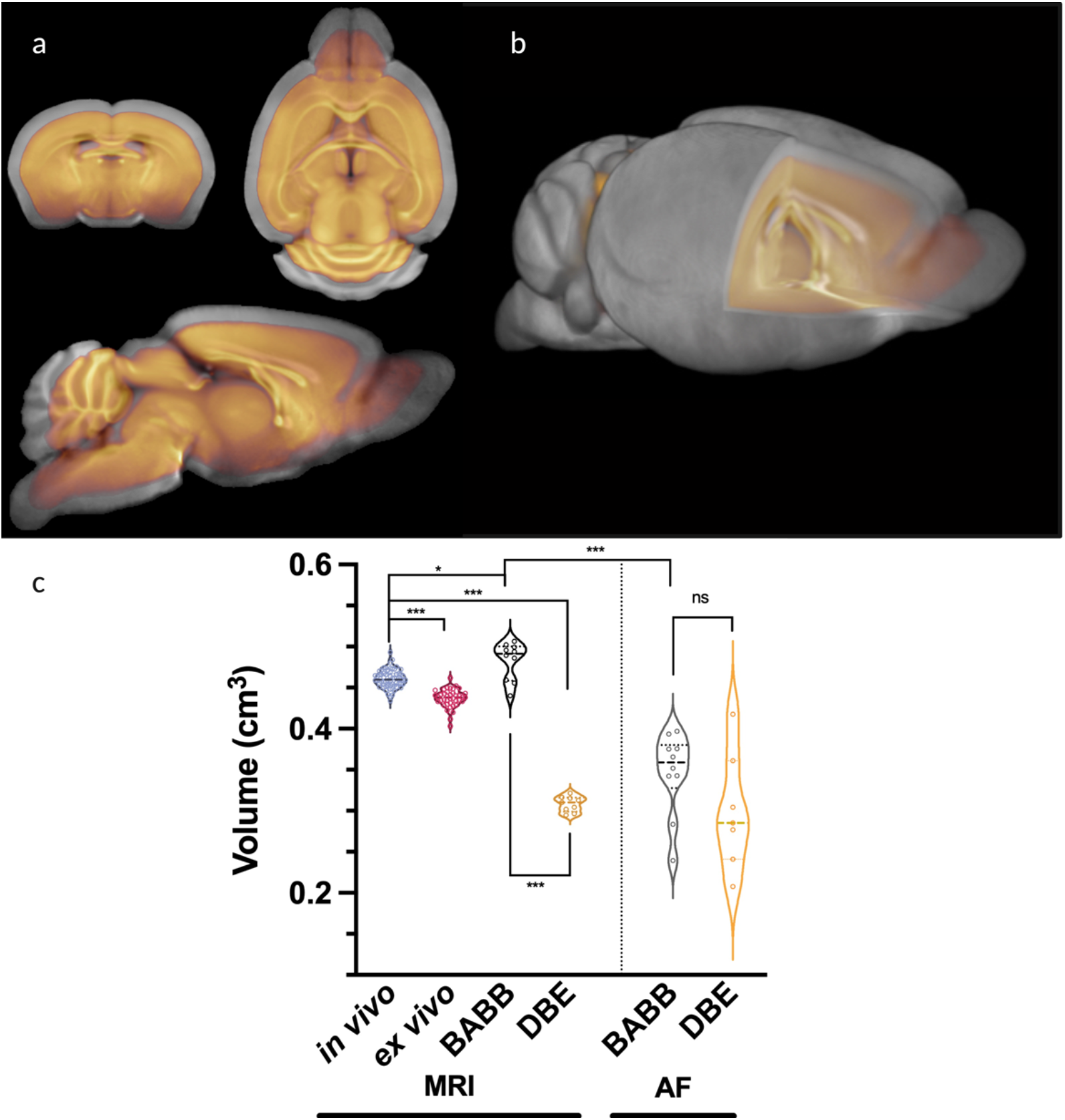
Differential effects of clearing methods on brain volume. a) Overlay of the average BABB template (n=10) (grey) and DBE template (n=9) (orange), indicating a clear difference in brain size. b) 3D overlay of the average BABB (grey) and DBE (orange) templates wherein the DBE template lies completely within the average BABB brain. c) Brain volume calculations of the average BABB and DBE brains with average *in vivo* and *ex vivo in situ* brain volumes, as well as with their respective autofluorescence volumes. All values are expressed in cm^3^. The average brain volume was significantly lower (***p<0.001) for DBE cleared brains (0.308 ± 0.009 cm^3^) as compared to BABB cleared brains (0.483 ± 0.023 cm^3^). The *in vivo* brain volume showed significant differences (***p<0.001; *p=0.03) with all other calculated brain volumes. Comparison of the average BABB and DBE brain sizes, based on autofluorescence, indicated no difference in brain size after optical clearing. Comparison of these volumes with their respective volumes after rehydration for MR-acquisition, indicated that the BABB brain volume was significantly larger when calculated based on T1-weighted images (0.483 ± 0.023 cm^3^) as when calculated from tissue (p<0.001) autofluorescence (0.343 ± 0.0517 cm^3^), which was not the case for the DBE cleared brains (p=0.94).

### OCUM: T1-weighted reference template and atlas for optically cleared brains

Due to the differential effects of BABB and DBE clearing and the significant size differences of the T1-weighted images (Figure 2), we decided on the creation of two brain templates containing all T1-weighted images (n=19) namely the Optically Cleared UMeå (OCUM) brain template and atlas for BABB/Murray’s cleared brains and the iOCUM brain template and atlas for DBE/iDISCO cleared brains. The DSURQE mouse brain template with 40 μm^3^ isotropic voxel size^14,21-24^ and its atlas labels served as the basis to construct both OCUM and iOCUM. A schematic overview of the applied pipeline to create OCUM and iOCUM is provided in Figure 3. All required image transformations were calculated in SPM, using the SPMmouse toolbox (http://spmmouse.org)^25^. The outcome of all applied transformations was reviewed cautiously by two independent readers and correctness and preservation of left and right was checked in every step to avoid involuntary image flipping along the y-axis. Initially, all individual BABB and DBE cleared brains were co-registered to their respective template (mid-point average of serial registration) with an estimation separation of 0.08 mm and 0.04 mm followed by 4^th^ degree B-spline interpolation. Co-registration was followed by normalization of the individual BABB brains to the DBE template and vice versa. These normalizations performed best with following settings: no affine regularization, trilinear interpolation and image smoothing using a Gaussian kernel of 0.72 mm. Of note, these specific normalizations performed worse when applying a higher degree B-spline transformation. Next, we created the OCUM and iOCUM templates by calculating the mid-point average from the serial registration of all individual (n=19), bias-corrected MR-images in BABB and DBE size, respectively (Figure 4a and 4b). Consequently, we created specific tissue segments and tissue probability maps (TPM) using a double consequent segmentation + DARTEL pipeline. For the initial segmentation and DARTEL pipeline, generating preliminary tissue priors for our templates, we used in-house tissue priors obtained from *ex vivo in situ* acquired T1-weighted images. Therefore, GM, WM and CSF segments were co-registered and normalized to both OCUM and iOCUM and applied as animal preset in SPMmouse. Thereafter, we ran the complete process (segmentation + DARTEL algorithm) again, using the preliminary tissue priors generated in the previous step, to produce accurate and template specific TPMs for both OCUM and iOCUM (Figure 4c). To use our newly created TPMs in all following image transformations and image analyses, we created two template-specific animal presets to use in the SPMmouse toolbox. Finally, we normalized (no affine regularization, 4^th^ degree B-spline interpolation and 16 non-linear iterations) the DSURQE atlas to OCUM space to delineate the ROIs. Subcortical structures such as the deep nuclei and the anterior cortical regions were perfectly aligned with OCUM by normalization and the posterior cortical ROIs were manually adapted on each template, resulting in OCUM and iOCUM specific atlas labels (Figure 4d). Together, our final resources (OCUM and iOCUM) comprise: two high-resolution (40 μm^3^ isotropic voxel dimension) whole mouse brain templates, their corresponding TPMs (GM + WM + CSF probability maps) required for brain segmentation and warping, a mouse brain atlas with 336 annotated ROIs and a protocol to create fluorescence-MR fusion images by means of co-registration and normalization.

**Figure 3:**
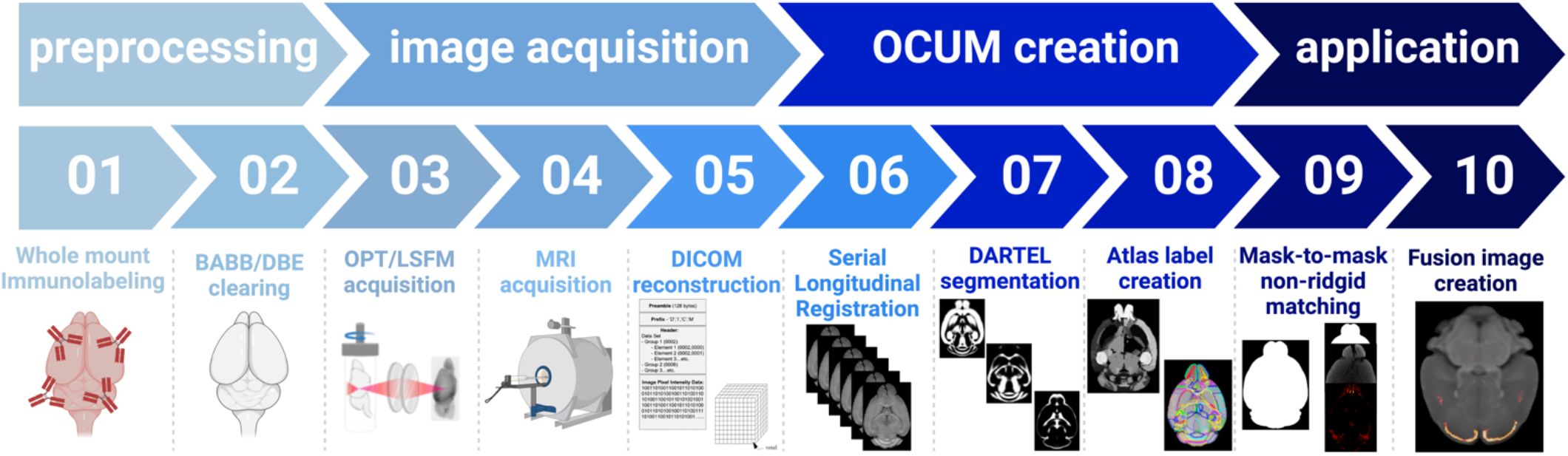
Schematic overview of the applied pipeline to obtain the OCUM brain template with its associated tissue priors and atlas labels.

**Figure 4:**
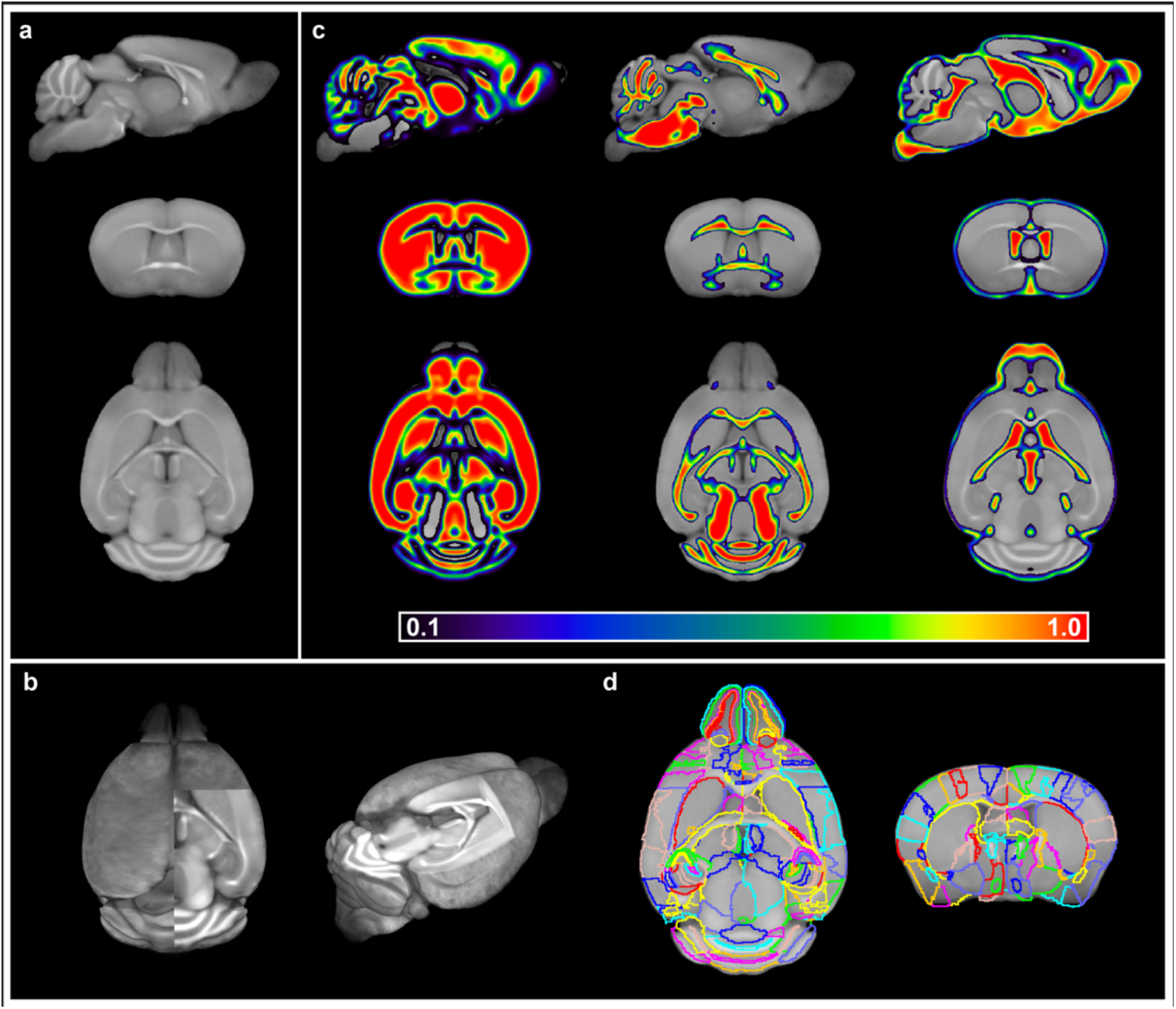
OCUM brain template and atlas. a) Sagittal, coronal and axial brain slice of the OCUM template (n=19). b) 3D representation of OCUM showing high GM and WM contrast in the template. c) GM, WM and CSF tissue probability maps associated with OCUM. d) Volume of interest (VOI) delineation exemplified in a sagittal and coronal slice of OCUM.

### Accurate anatomical referencing of optical brain images

To highlight the applicability of the newly designed templates, we created fusion images of fluorescent optical signals, acquired from BABB and DBE cleared brains, with OCUM and iOCUM, respectively. Therefore, optical images were reconstructed into DICOM format and converted to NIFTI format (Figure 3), to allow image transformations in SPM and fusion image creation in PMOD. First, we co-registered (voxel-to-voxel affine registration, followed by 4^th^ degree B-spline interpolation) the optical images to the template. Therefore, voxel-to-voxel affine transformation matrices were calculated using the autofluorescence image. Since the autofluorescence and signal images are in the same native space, this transformation matrix can then be applied to the signal image to co-register fluorescent signals to the template. To improve quality of the fusion images by compensating for natural variation in brain size and differences in tissue deformation due to dehydration and clearing, normalization or warping to the template image is preferred. As shown in figure 1, anatomical detail remains limited in the autofluorescence image, which hampers accurate image warping. Therefore, we created a binary mask based on the autofluorescence image and normalized (no affine regularization, nearest-neighbor interpolation, and 2 mm gaussian kernel smoothing) this mask to a similar binary (i)OCUM mask and applied identical transformations to the original images which resulted in near-perfect normalized fusion images (Figure 5, Supplementary Figure 2). In each case, fusion images allowed for precise anatomical identification of brain regions with fluorescent signal. In figure 5a, we traced anterograde innervation from the eye after injection of fluorescently labeled cholera toxin B (CTB) in the anterior chamber of the eye by visualizing CTB using OPT. We observed clear localization of CTB signal within LGN of thalamus and the superior colliculus and observed a perfect match of brain regions of the visual system, indicating anterograde innervation from the eye towards the visual cortex. Figure 5b and supplementary figure 2 display fusion images from DAT OPT with OCUM and iOCUM, respectively. We identified clear DAT signal in the striatum, hypothalamus, olfactory tubercle, amygdala and even in the substantia nigra. All these regions are well-known to express DAT, which underlines the applicability of our templates and atlases. Lastly, we created fusion images of both OPT (figure 5c) and LSFM (figure 5d) signal of TSPO, a well-known microglial neuroinflammation marker, with OCUM. In contrast to the two previous examples, TSPO displays a diffuse expression pattern rather than being expressed in distinct brain regions. Figures 5c and d clearly show that the presented pipeline also works for optical images originating from diffusely expressed markers. Both TSPO OPT and LSFM showed elevated signal in brain vasculature, which can be explained by TSPO expression in endothelial cells. Furthermore, next to detailed vessel staining, TSPO LSFM showed high signal in the cortical layer IV, which suggests higher expression levels of activated microglia in this highly myelinated cortical layer.

**Figure 5:**
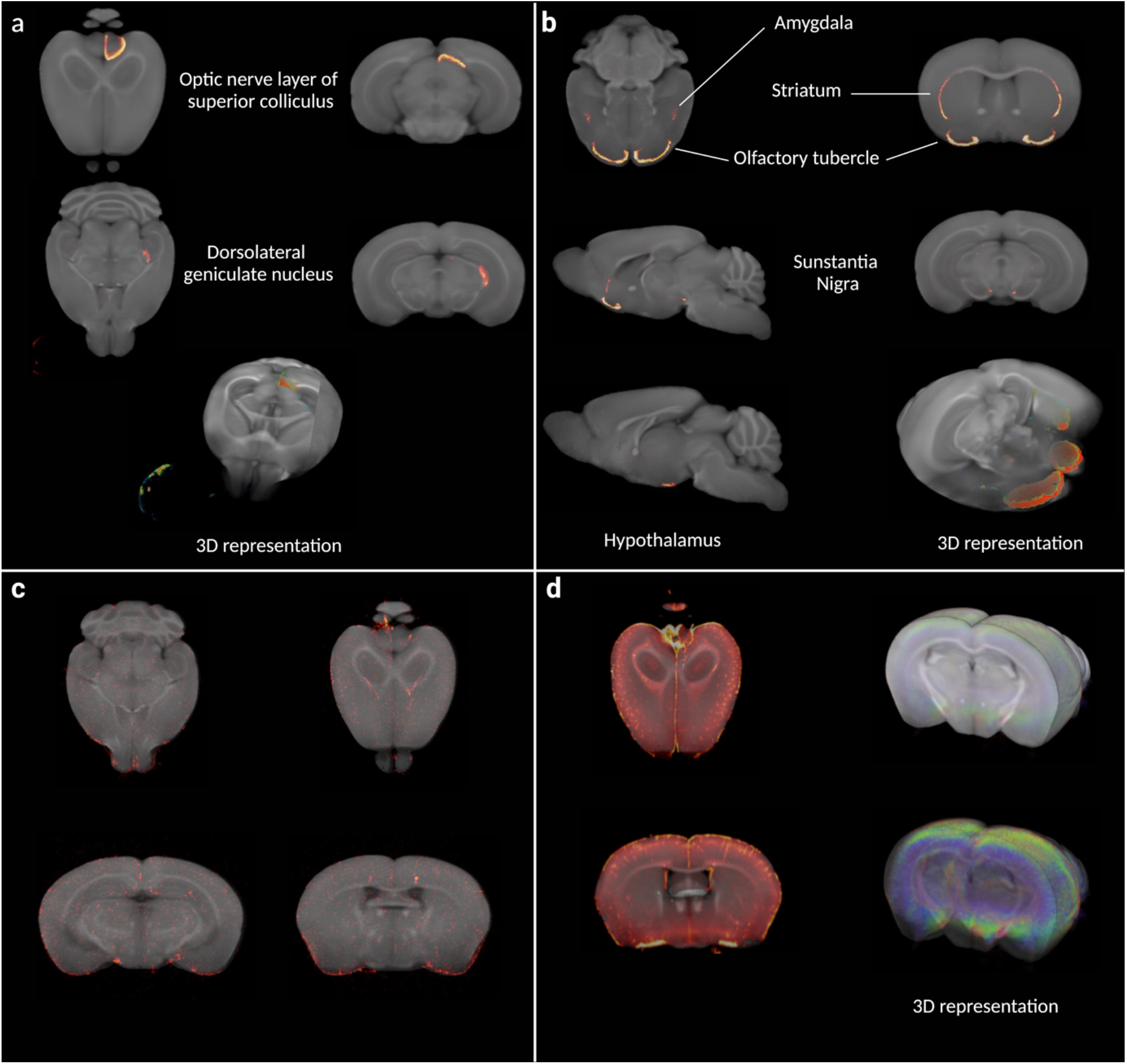
Fusion images of 3D optical signal with the OCUM template. a) Fusion images of OPT signal of cholera toxin B with OCUM displaying clear optical signal in distinct parts of the visual system after intraocular injection. b) Fusion images of OPT signal from typical dopamine transporter expressing brain regions and OCUM. c) Fusion images of OPT signal from TSPO, targeting microglia, and OCUM. d) Fusion images of LSFM signal from TSPO and OCUM showing high optical signal in cortical layer 4 and intense vessel staining due to TSPO expression in endothelial cells.

## Discussion

Whole brain optical imaging is rapidly gaining interest and popularity for the study of protein expression profiles and disease markers in neuroscience. MR, on the other hand, is a non-invasive, well-established and common imaging modality used for brain research, both in preclinical and clinical settings. Application of these techniques, in combination with advanced image quantification, represents a powerful triad to discover insights into both the healthy and diseased brain. Here we report, to our knowledge, the first MR-based high-resolution brain template and atlas, specifically designed for brains that were subjected to the required pre-processing and tissue clearing protocols for 3D optical imaging. To overcome issues related to differential volumetric and morphological effects related to tissue clearing, two versions of the template were created: OCUM for BABB/Murray’s clearing method and iOCUM for DBE/iDISCO protocols. The utility and application of both templates was then illustrated by detailed anatomical mapping of several distinct whole brain optical signals.

To optimally design OCUM and iOCUM as MRI-based template atlases, we employed the DSURQE template and atlas^14,21-24^ as a starting point. While the AIBS CCFv3 has been used as a reference for brain atlases in several publications involving LSFM^10,11^, we specifically chose to work with an MRI-based template as a starting point, rather than a two-photon tomography-based template, to increase accuracy. Our resulting resources are comprised of four key components namely: (1) a T1-weighted template image (40 μm^3^ isotropic spatial resolution) defining the (i)OCUM space; (2) the associated tissue priors, or TPMs, required to segment and normalize optical images to the template space; (3) a whole brain atlas, delineating 336 ROIs and (4) a detailed protocol of how to employ these resources to create fusion images and identify specific areas containing optical signal using the atlas. These resources will furthermore allow for ROI- and voxel-based quantitative analysis of large samples of brains, aligned in the same image space, both being well-characterized quantification methods for PET and MR brain studies. It should be noted that each template is comprised of brain images acquired from both C57Bl/6J WT and transgenic mice, however, all transgenic models were bred on a C57Bl/6 background. The knockout mice used to create the templates did not have any differences in brain size or morphology to C57Bl/6J WT mice, and therefore did not influence the anatomical precision of either template. Since (i)OCUM is based on normal adult C57Bl/6 mouse brains, its application might not be justified when using mice with severe brain defects or altered brain morphology and optical data obtained from other mouse strains must be cautiously handled.

The fact that different tissue clearing methods exert differential effects on brain size and morphology has become a generally accepted concept in the field^16,26^. Our head-to-head comparison showed significant volume differences calculated from T1-weighted images between BABB and DBE-cleared brains (Figure 2), while no significant differences in brain volume were detected when calculated using tissue autofluorescence (Figure 2c). Interestingly, there was no significant difference in DBE cleared brain volume calculated from autofluorescence or T1-weighted images, while for BABB cleared brains, volumes calculated from T1-weighted images was even slightly larger than *in vivo* brain volume. These data suggest a differential effect of clearing agents on tissue rehydration rather than on shrinkage. Another factor we identified to impact brain size of optical images, potentially rendering the pipeline more challenging, was the zoom factor used during whole brain imaging. With our OPT setup, we repeatedly observed near perfect fusion results on images acquired with an optical zoom factor of 1.25 and 1.6 during image acquisition, while images acquired with larger zoom factors were less accurate after normalization, likely due to additional skewing to OCUM. In line with previous reports mapping optical 3D signal to brain templates^10^, we experienced more difficulty in automatically delineating the cortical ROIs located in the hind brain and had to adapt them manually to fit, while this procedure was not required in anterior cortical regions, implying differential effects of clearing agents throughout the brain.

The creation of optical-MR fusion images by means of co-registration and normalization implies bringing the optical signal to the MR template reference space, adapting these both to the template’s bounding box and voxel dimension. This has several implications for optical images. While OCUM is a high-resolution template with a voxel size of 40 μm^3^, the original OPT-images in this study have a voxel size ranging from 16.5 – 21 μm^3^. This means that the voxel size of optical images is increased, thus lowering their resolution to create fusion images to perform anatomical mapping. LSFM has even smaller voxel dimensions, hence higher resolution compared to OPT, resulting in a greater loss of resolution when fit into the MR-template space. Furthermore, while OPT imaging generates isotropic voxels (identical dimensions along x, y, and z-axis), LSFM has lower axial than lateral resolution, resulting in anisotropic voxels. LSFM images can be first resampled along the z-axis to reach isotropic voxel size and then co-registered and normalized to the anatomical template. Nevertheless, this may lead to ambiguities in the axial direction due to signal skewing during the resampling process, which may greatly impact voxel-wise quantitative analyses but may even exert a clear effect on the ROI level. However, great advances are being made both for software and hardware in this field. In 2019, Chakraborty et al. described cleared-tissue axially swept light-sheet microscopy (ctASLM) wherein z-axial resolution is significantly increased which results in approximate isotropic voxels^27^.

To exemplify the possibilities of our resources, we created fusion images of OPT and LSFM images with OCUM and iOCUM, for BABB and DBE clearing, respectively. Using both OCUM and iOCUM, we created fusion images of DAT expression in the mouse brain and identified signal in the striatum, amygdala, olfactory tubercle, hypothalamus and substantia nigra. All these regions are known DAT expressing regions, which clearly highlight the applicability of our resources. Although beyond the scope of this study, we only observed DAT signal on the outermost surface of striatum based on OPT, which might be due to the lower resolution of OPT compared to LSFM or due to antibody competition induced by the high striatal expression rate of the transporter. Furthermore, we were able to trace anterograde innervation from the eye to LGN of thalamus and superior colliculus and observed a perfect match to brain regions from the visual system by the optic nerve. Finally, we visualized TSPO expression both by OPT and LSFM and observed high expression on the 4^th^ cortical brain layer indicating high levels of microglia in this layer and observed extremely high signal in the vessels. Although it is known that TSPO is also expressed in endothelial cells^28^, TSPO Positron Emission Tomography Images do not show this feature due to its limited resolution while OPT and LSFM showed the extent of vessel binding of this important neuroinflammation marker. These examples highlight the potential of this technique to discover novel biological insights among different brain systems and brain diseases. Indeed, OCUM was recently used to identify viral distribution patterns on a whole brain level using OPT^29^. Together, this demonstrates how these resources can aid spatial and quantitative analyses of treated versus control animals or for a cross-sectional quantitation of specific disease markers over time. Finally, it may serve in the further development of machine-learning approaches for optical imaging^30^.

In conclusion, we present a full MRI-based brain template and atlas for mouse brains that were previously processed and cleared for 3D whole brain optical imaging. Thereby, we provide the brain imaging community with a unique tool allowing anatomical brain mapping of optical brain signals without the need for repetitive, time-consuming, and expensive MRI scanning. Furthermore, OCUM and iOCUM offer a means to standardize structural and functional optical data analysis pipelines that may significantly assist in the discovery of novel neurobiological insights.

## Methods

### Ethics declaration

All animal experiments were approved and performed according to the guidelines of the regional Animal Research Ethics Committee of Northern Norrland, the Animal Review Board at the Court of Appeal of Northern Stockholm and by the Swedish Board of Agriculture (Ethical permits: A35-2016, A9-2018 and A41-2019). Reporting regarding all *in vivo* experiments was performed compliant with the ARRIVE guidelines.

### Animals

Eight- to eleven-week-old male C57Bl/6J mice (n=10) were purchased from Jackson Laboratories (Bar Harbor, ME, USA) or and Charles River, (Wilmington, MA, USA). Interferon alpha/beta receptor knockout (IFNAR^-/-^) (n=6, 4M/2F) (Muller 1994), interferon-beta promoter stimulator-1 knockout (IPS-1^-/-^) (n=3, 1M/2F) and Viperin^-/-^(n=1, F) mice (a kind gift from Peter Cresswell, Department of Immunobiology, Yale University School of Medicine) were bred at the Umeå Centre for Comparative Biology (UCCB). Animal experiments were conducted at UCCB and at the department of Molecular Medicine and Surgery at Karolinska Institutet (MMK). Following euthanasia using O_2_ deprivation or anesthesia using 60 mg/ml pentobarbital (APL, Kungens Kurva, Sweden), all animals (n=19) were transcardially perfused using 20 mL PBS followed by 20 mL 4% paraformaldehyde (PFA) in PBS whereafter brains were harvested for *ex vivo* analyses.

### Whole *mount immunohistochemistry and optical clearing*

PFA-fixed brains were fluorescently immunolabeled and processed for OPT as described previously^3,31^. Briefly, brains were dehydrated using stepwise gradients of methanol (MeOH), permeabilized by 4 cycles of repetitive freeze-thawing in MeOH at -80°C and bleached overnight (ON) in MeOH:H_2_O_2_:DMSO (2:3:1) at room temperature (RT) to quench tissue autofluorescence. For immunolabeling, brains were rehydrated into TBST (50mM Tris-HCl pH7.4, 150mM NaCl and 0.1% TritonX-100) and labelled with primary (recombinant rabbit anti-TSPO (1:1000) (ab109497, Abcam, Camebridge, United Kingdom) or rabbit anti-DAT (1:400) (clone 1D2 ZooMAb, n°: ZRB1525, Sigma Aldrich, St. Louis, MO, USA) and secondary (goat anti-rabbit Alexa-594 (1:500) (A-11037, Thermo Fisher, Scientific, Waltham, MA, USA) or donkey anti-rabbit Alexa-594 (1:500) (ab150064, Abcam)). After immunolabeling, all brains were mounted in 1.5% low melting point agarose (SeaPlaque, Lonza, Basel, Switzerland) and optically cleared using benzyl alcohol: benzyl benzoate (1:2) (BABB) or dibenzyl ether (DBE) (Sigma-Aldrich).

### Optical Projection Tomography

OPT image acquisition was performed on an in-house developed near-infrared OPT (NiR-OPT) system, as described by Eriksson et al.^31^ A zoom factor of 1.25 (cholera toxin) or 1.6 (all other brains) was applied which resulted in a respective isotropic voxel dimension of 21 μm^3^ or 16.5 μm^3^. For cholera toxin, OPT images were acquired using the following settings: Ex: 630/50 nm, Em: 665/95nm (exposure time: 3000 ms). For all other targets, OPT images were acquired using Ex: 580/25 nm, Em: 625/30 nm (exposure time: 500 ms (TSPO) or 3000 ms (DAT). All tissue fluorescence images were acquired with the same settings namely: Ex: 425/60 nm, Em 480LP nm (exposure time: 200 ms). To increase the signal-to-noise ratio (SNR) of the labeled molecules in the brains, the pixel intensity range of all images were adjusted to display minima and maxima and a contrast limited adaptive histogram equalization (CLAHE) algorithm with a tile size of 16 × 16 was applied to the projection images acquired in the fluorescent signal channels. Tomographic reconstruction with additional misalignment compensation and ring artifact reduction was performed using NRecon software v.1.7.0.4 (Skyscan microCT, Bruker, Belgium). Afterwards, OPT images displaying both the targeted signals and the tissue autofluorescence signals, were reconstructed into DICOM format using NRecon software, followed by their conversion into NifTi format using the PMOD view tool (version 4.2, PMOD Technologies Inc., Zurich, Switzerland).

### Light Sheet Fluorescence Microscopy

The brain stained for TSPO and imaged by NiR-OPT was consequently rescanned using an UltraMicroscope II (Miltenyi Biotec, Germany) including a 1x Olympus objective (Olympus PLAPO 2XC) coupled to an Olympus MVX10 zoom body, providing 0.63x up to 6.3x magnification with a lens corrected dipping cap MVPLAPO 2x DC DBE objective (Olympus). The cleared brain was immerged in BABB and magnification was set to 0.63x. For image acquisition, left and right light sheets were blend merged with a 0.14 numerical aperture, resulting in a light sheet z-thickness of 3.87μm and 80% width, while using a 12-step contrast adaptive dynamic focus across the field of view. Image sections with a step size of 10 μm were generated by Imspector Pro software (v7.1.15, Miltenyi Biotec Gmbh, Germany) and stitched together using the implemented TeraStitcher script (v.9). The obtained images were then converted into NifTi files using Amira Avizo software (version 6.3.0, Thermo Fisher Scientific, Waltham, MA, USA) and resampled prior to coregistration.

### MRI acquisition

After optical clearing with BABB (n=10) or DBE (n=9) and OPT scanning of selected brains, all brains (n=19), were rehydrated into TBST, incubated in 0.29M sucrose to remove the surrounding agarose and washed in PBS prior to MRI. T1-weightd images were acquired using a Modified Driven Equilibrium Fourier Transform (MDEFT) sequence with five repetitions (TR: 3000 ms; TE: 3 ms; TI: 950 ms; voxel size 40 × 40 × 40 μm^3^) on a 9.4 Tesla (T) preclinical MR system (Bruker BioSpec 94/20, Bruker Ettlingen, Germany), equipped with a cryogenic Radio Frequency (RF) coil (MRI CryoProbe, Bruker) running Paravision 7.0 software. Data were exported in DICOM format using Paravision routines followed by image conversion from DICOM to NifTi format using the dcm2nii tool in MRIcron. The individual repetitions of each scan were realigned and averaged using statistical parametric mapping (SPM8) (the Wellcone Trust Centre for Neuroimaging, UCL, London, U.K.) implemented in Matlab (R2014a, The MathWorks Inc., Natick, MA, USA).

### OCUM template and atlas creation

Initially, two distinct templates: namely one specifically for BABB (n=10) and DBE (n=9) cleared brains, were created using bias corrected (SPM8) MR images, which were realigned and averages using serial longitudinal registration (SLR) in SPM12, implemented in Matlab (R2015b, The MathWorks Inc.) whereafter all individual MR scans were coregistered to their respective template. Individual DBE brains were then normalized to the BABB template while individual BABB brains were normalized to the DBE template. Consequently, the final OCUM and iOCUM templates were created by rerunning the SLR on all brains (n=19) both in BABB and DBE size, respectively. Both for OCUM and iOCUM, specific segments and TPMs were created using a 2-step segmentation and DARTEL pipeline, initially based on in-house generated tissue priors. Briefly, a primary segmentation and DARTEL algorithm was applied to the individual MR images of both templates to generate preliminary tissue priors for both OCUM and iOCUM, using the toolbox SPMmouse^25^. Thereafter, the complete process (segmentation + DARTEL) was repeated using the tissue priors generated from the previous step to produce accurate template specific TPMs for both templates.

### Creation of fusion images with OCUM template

Initially, both the autofluorescence image and the OPT image displaying specific signals were reoriented manually in SPM8 to the templates orientation and their origins were set tangent to the upper edge of the brain at Bregma. For co-registration of OPT and MR images, voxel-to-voxel affine transformation matrices were calculated using the autofluorescence OPT images and applied to those displaying the specifically targeted signals. To further improve the fusion images and enable image warping, binary masks of the autofluorescence OPT images were created in ITK-SNAP version 3.8.0 (www.itksnap.org)^32^ which were consequently normalized to a binary OCUM template mask. Finally, fusion images were created in the PMOD view tool and 3D images were created in Amira-Avizo software (version 6.3.0, Thermo Fisher Scientific).

## Data availability statement

All data are available at the department of Clinical Microbiology of Umeå University and can be obtained upon request. The OCUM and iOCUM brain templates with all their resources will be made publicly available for general use upon acceptance of the manuscript.

## Acknowledgements

The research leading to this publication has received funding from the Umeå University Medical Faculty (D.M. and U.A.), the Kempe Foundations (U.A.), the Swedish Research Council (A.K.Ö. and P.O.B), the Laboratory for Molecular Infection Medicine Sweden (MIMS) (A.K.Ö), the Novo Nordisk Foundation (M.V. and P.O.B) and the Family Erling Persson Foundation (P.O.B). S.M.A.W, is the holder of a Marie Curie Actions postdoctoral fellowship funded by the European Commission. N.C is the holder of a stipend from the MIMS Excellence by Choice Postdoctoral Program, funded by the Knut and Alice Wallenberg Foundation (KAW2015.0284). We thank the Small Animal Research and Imaging Facility (SARIF) at Umeå University for providing equipment and technical support for performing parts of the study. We thank Dr. C. Nord for technical support in advanced image reconstruction.

## Additional information

Supplementary information accompanies this paper.

The authors declare that there are no competing interests.

## Figures and Figure legends

**Supplementary Figure 1:**
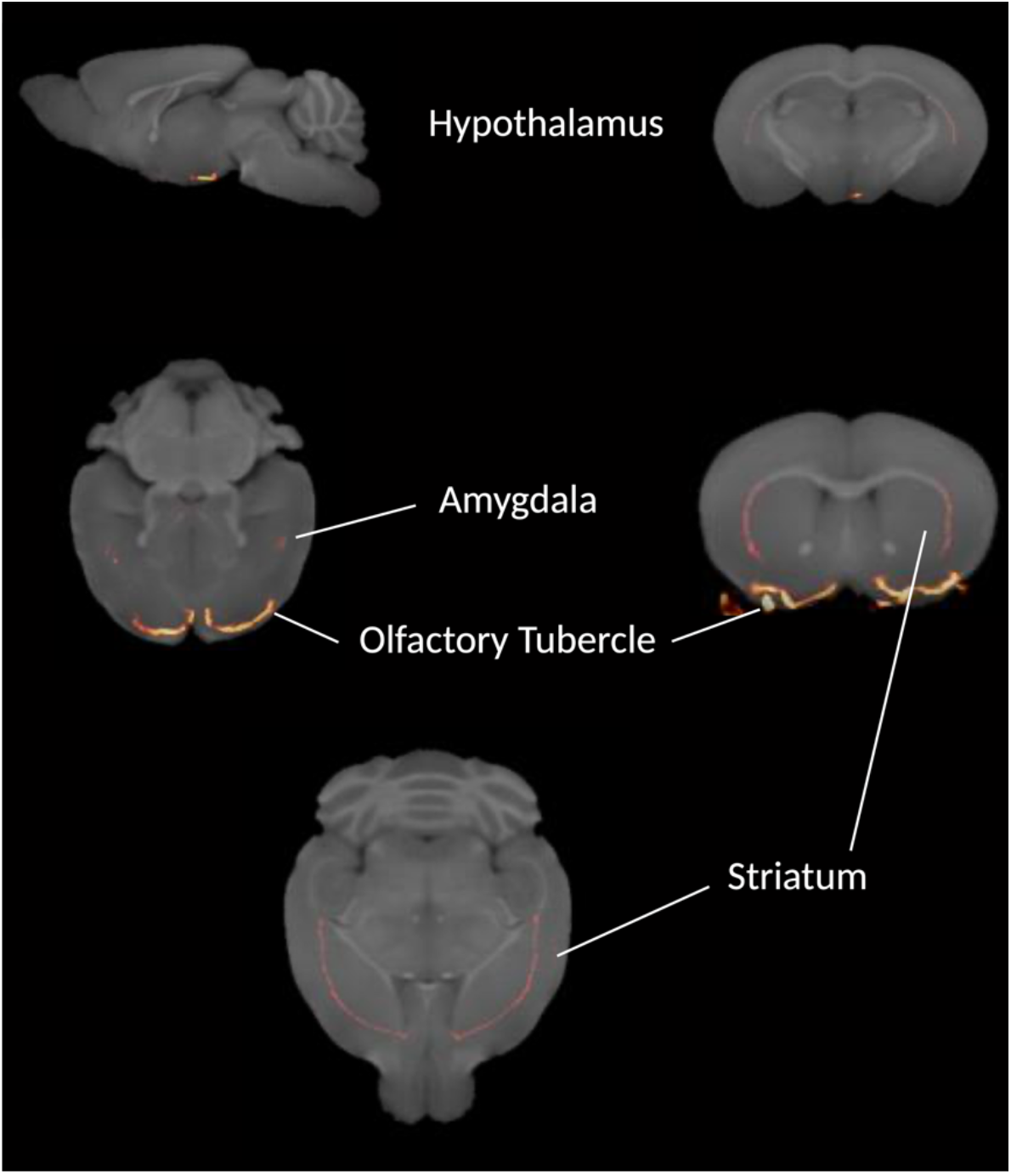
Fusion images of 3D optical signal with the iOCUM template. Fusion images of OPT signal from typical dopamine transporter expressing brain regions in DBE cleared brains with iOCUM.

**Supplementary Table 1:**
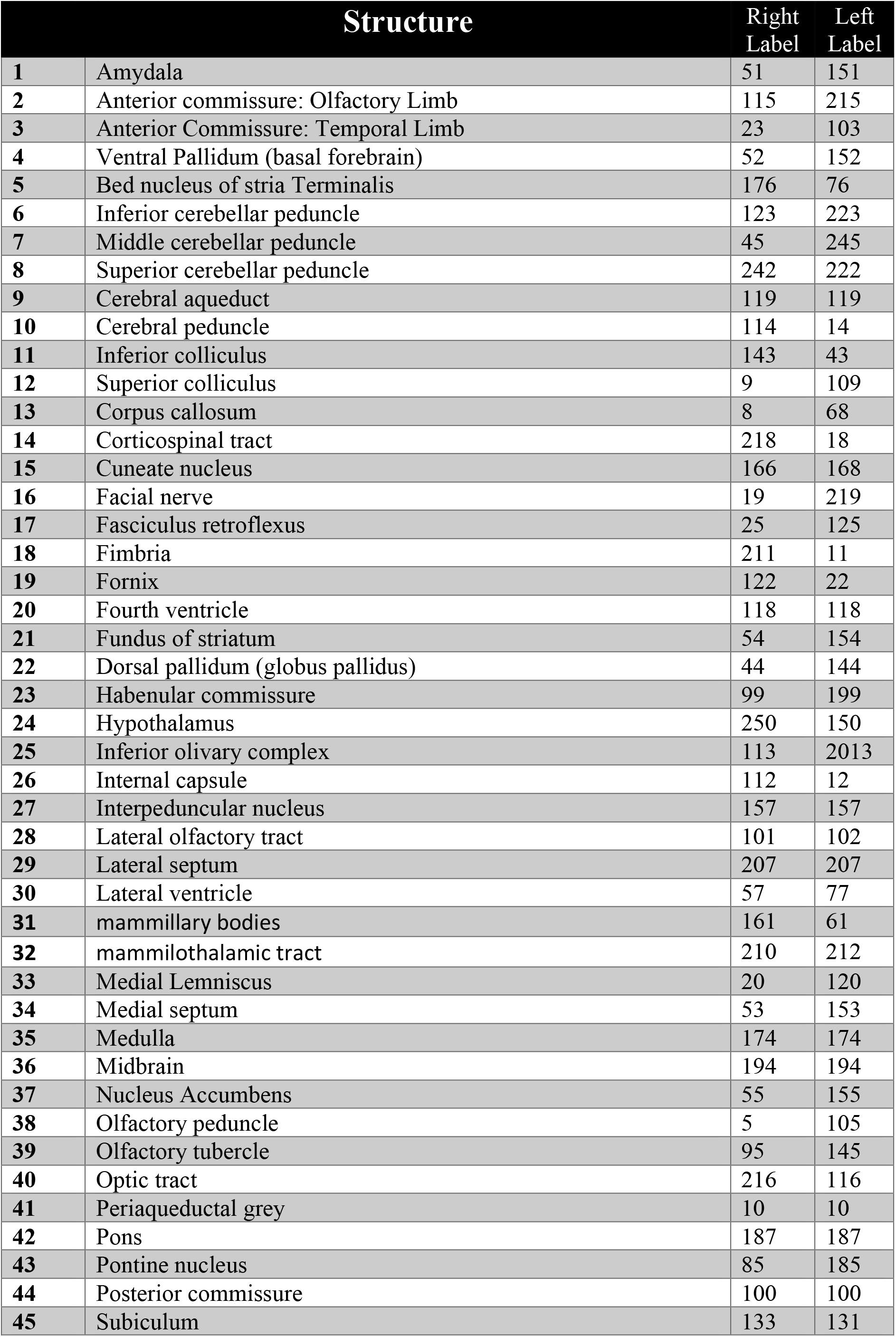
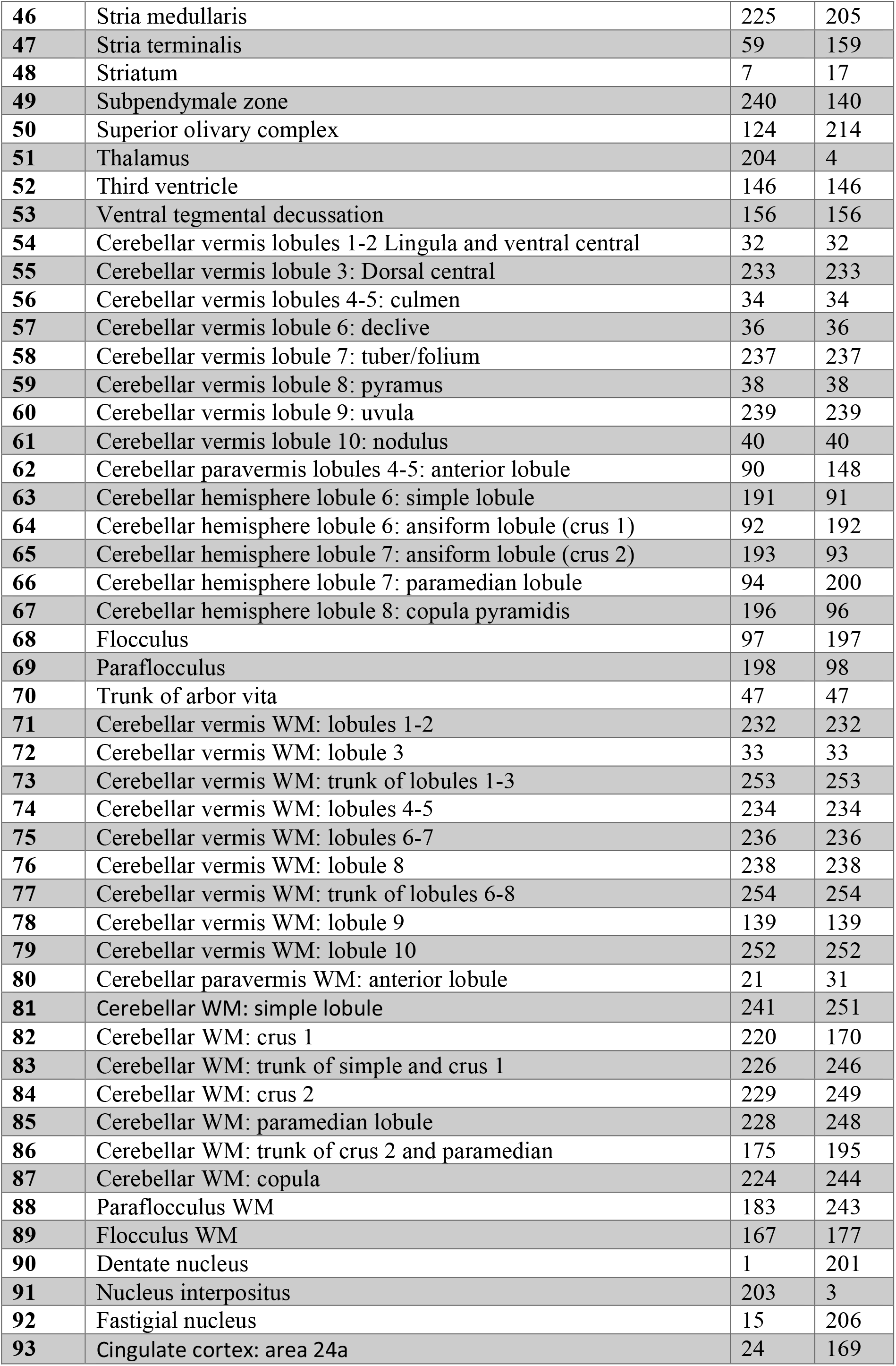
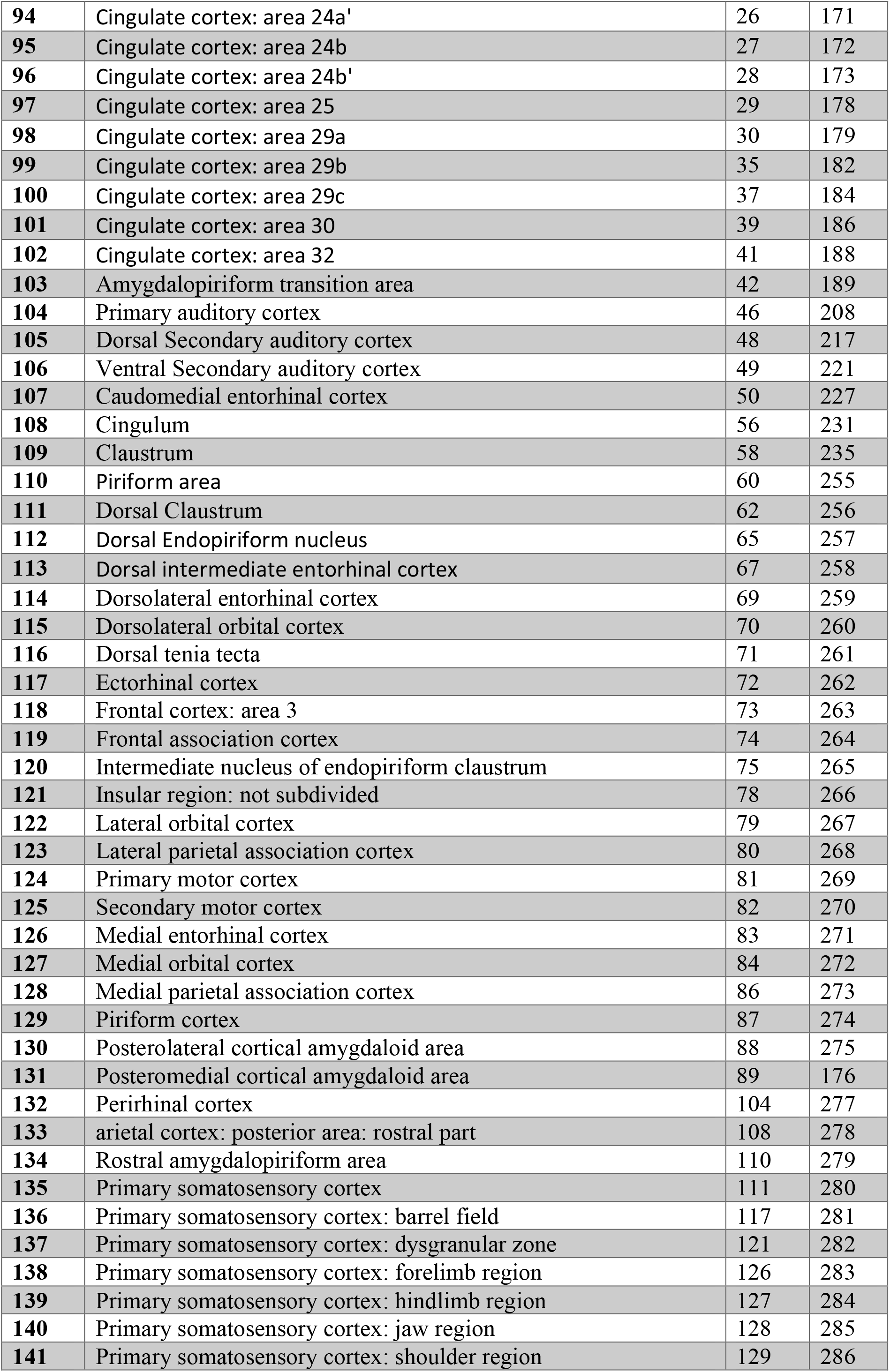
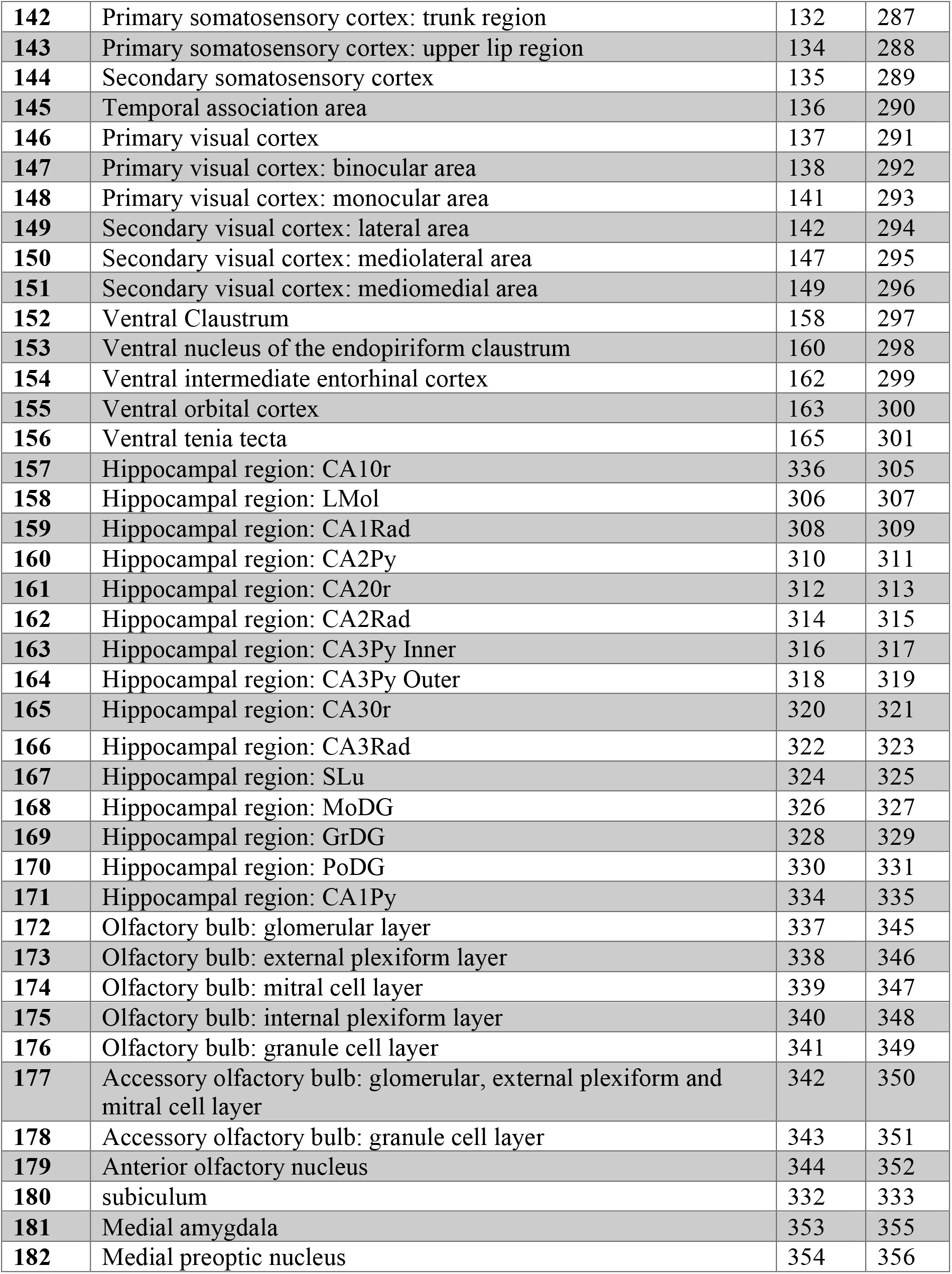
OCUM and iOCUM atlas labels.

## Notes

### Competing Interest Statement

The authors have declared no competing interest.

## References

1 Dodt, H. U. et al. Ultramicroscopy: three-dimensional visualization of neuronal networks in the whole mouse brain. Nat Methods 4, 331–336, doi:10.1038/nmeth1036 (2007).

2 Sharpe, J. et al. Optical projection tomography as a tool for 3D microscopy and gene expression studies. Science 296, 541–545, doi:10.1126/science.1068206 (2002).

3 Alanentalo, T. et al. Tomographic molecular imaging and 3D quantification within adult mouse organs. Nat Methods 4, 31–33, doi:10.1038/nmeth985 (2007).

4 Hansen, H. H., Roostalu, U. & Hecksher-Sorensen, J. Whole-brain three-dimensional imaging for quantification of drug targets and treatment effects in mouse models of neurodegenerative diseases. Neural Regen Res 15, 2255–2257, doi:10.4103/1673-5374.284983 (2020).

5 Wang, Q. et al. The Allen Mouse Brain Common Coordinate Framework: A 3D Reference Atlas. Cell 181, 936–953 e920, doi:10.1016/j.cell.2020.04.007 (2020).

6 Oh, S. W. et al. A mesoscale connectome of the mouse brain. Nature 508, 207–214, doi:10.1038/nature13186 (2014).

7 Sunkin, S. M. et al. Allen Brain Atlas: an integrated spatio-temporal portal for exploring the central nervous system. Nucleic Acids Res 41, D996–D1008, doi:10.1093/nar/gks1042 (2013).

8 Furth, D. et al. An interactive framework for whole-brain maps at cellular resolution. Nat Neurosci 21, 139–149, doi:10.1038/s41593-017-0027-7 (2018).

9 Renier, N. et al. Mapping of Brain Activity by Automated Volume Analysis of Immediate Early Genes. Cell 165, 1789–1802, doi:10.1016/j.cell.2016.05.007 (2016).

10 Perens, J. et al. An Optimized Mouse Brain Atlas for Automated Mapping and Quantification of Neuronal Activity Using iDISCO+ and Light Sheet Fluorescence Microscopy. Neuroinformatics 19, 433–446, doi:10.1007/s12021-020-09490-8 (2021).

11 Salinas, C. B. G. et al. Integrated Brain Atlas for Unbiased Mapping of Nervous System Effects Following Liraglutide Treatment. Sci Rep 8, 10310, doi:10.1038/s41598-018-28496-6 (2018).

12 Qu, L. et al. Cross-modal coherent registration of whole mouse brains. Nat Methods 19, 111–118, doi:10.1038/s41592-021-01334-w (2022).

13 Barriere, D. A. et al. Brain orchestration of pregnancy and maternal behavior in mice: A longitudinal morphometric study. Neuroimage 230, 117776, doi:10.1016/j.neuroimage.2021.117776 (2021).

14 Dorr, A. E., Lerch, J. P., Spring, S., Kabani, N. & Henkelman, R. M. High resolution three-dimensional brain atlas using an average magnetic resonance image of 40 adult C57Bl/6J mice. Neuroimage 42, 60–69, doi:10.1016/j.neuroimage.2008.03.037 (2008).

15 Hikishima, K. et al. In vivo microscopic voxel-based morphometry with a brain template to characterize strain-specific structures in the mouse brain. Sci Rep 7, 85, doi:10.1038/s41598-017-00148-1 (2017).

16 Wan, P. et al. Evaluation of seven optical clearing methods in mouse brain. Neurophotonics 5, 035007, doi:10.1117/1.NPh.5.3.035007 (2018).

17 Hahn, M. et al. Mesoscopic 3D imaging of pancreatic cancer and Langerhans islets based on tissue autofluorescence. Sci Rep 10, 18246, doi:10.1038/s41598-020-74616-6 (2020).

18 Becker, K., Jahrling, N., Saghafi, S. & Dodt, H. U. Ultramicroscopy: light-sheet-based microscopy for imaging centimeter-sized objects with micrometer resolution. Cold Spring Harb Protoc 2013, 704–713, doi:10.1101/pdb.top076539 (2013).

19 Renier, N. et al. iDISCO: a simple, rapid method to immunolabel large tissue samples for volume imaging. Cell 159, 896–910, doi:10.1016/j.cell.2014.10.010 (2014).

20 Mediavilla, T. et al. Learning-related contraction of grey matter in rodent sensorimotor cortex is associated with adaptive myelination. Elife 11, doi:10.7554/eLife.77432 (2022).

21 Qiu, L. R. et al. Mouse MRI shows brain areas relatively larger in males emerge before those larger in females. Nat Commun 9, 2615, doi:10.1038/s41467-018-04921-2 (2018).

22 Richards, K. et al. Segmentation of the mouse hippocampal formation in magnetic resonance images. Neuroimage 58, 732–740, doi:10.1016/j.neuroimage.2011.06.025 (2011).

23 Steadman, P. E. et al. Genetic effects on cerebellar structure across mouse models of autism using a magnetic resonance imaging atlas. Autism Res 7, 124–137, doi:10.1002/aur.1344 (2014).

24 Ullmann, J. F., Watson, C., Janke, A. L., Kurniawan, N. D. & Reutens, D. C. A segmentation protocol and MRI atlas of the C57BL/6J mouse neocortex. Neuroimage 78, 196–203, doi:10.1016/j.neuroimage.2013.04.008 (2013).

25 Sawiak, S. J., Picq, J. L. & Dhenain, M. Voxel-based morphometry analyses of in vivo MRI in the aging mouse lemur primate. Front Aging Neurosci 6, 82, doi:10.3389/fnagi.2014.00082 (2014).

26 Kim, J. H. et al. Optimizing tissue-clearing conditions based on analysis of the critical factors affecting tissue-clearing procedures. Sci Rep 8, 12815, doi:10.1038/s41598-018-31153-7 (2018).

27 Chakraborty, T. et al. Light-sheet microscopy of cleared tissues with isotropic, subcellular resolution. Nat Methods 16, 1109–1113, doi:10.1038/s41592-019-0615-4 (2019).

28 Guilarte, T. R., Rodichkin, A. N., McGlothan, J. L., Acanda De La Rocha, A. M. & Azzam, D. J. Imaging neuroinflammation with TSPO: A new perspective on the cellular sources and subcellular localization. Pharmacol Ther 234, 108048, doi:10.1016/j.pharmthera.2021.108048 (2022).

29 Chotiwan, N. e. a. in bioRxiv (2021).

30 Todorov, M. I. et al. Machine learning analysis of whole mouse brain vasculature. Nat Methods 17, 442–449, doi:10.1038/s41592-020-0792-1 (2020).

31 Eriksson, A. U. et al. Near infrared optical projection tomography for assessments of beta-cell mass distribution in diabetes research. J Vis Exp, e50238, doi:10.3791/50238 (2013).

32 Yushkevich, P. A. et al. User-guided 3D active contour segmentation of anatomical structures: significantly improved efficiency and reliability. Neuroimage 31, 1116–1128, doi:10.1016/j.neuroimage.2006.01.015 (2006).

