## Supplementarty Table 1 for "An MR-based brain template and atlas for optical projection tomography and light sheet fluorescence microscopy"

Supplementary Table 1: OCUM and iOCUM atlas labels

|  | Structure | Right Label | Left Label |
| --- | --- | --- | --- |
| 1 | Amydala | 51 | 151 |
| 2 | Anterior commissure: Olfactory Limb | 115 | 215 |
| 3 | Anterior Commissure: Temporal Limb | 23 | 103 |
| 4 | Ventral Pallidum (basal forebrain) | 52 | 152 |
| 5 | Bed nucleus of stria Terminalis | 176 | 76 |
| 6 | Inferior cerebellar peduncle | 123 | 223 |
| 7 | Middle cerebellar peduncle | 45 | 245 |
| 8 | Superior cerebellar peduncle | 242 | 222 |
| 9 | Cerebral aqueduct | 119 | 119 |
| 10 | Cerebral peduncle | 114 | 14 |
| 11 | Inferior colliculus | 143 | 43 |
| 12 | Superior colliculus | 9 | 109 |
| 13 | Corpus callosum | 8 | 68 |
| 14 | Corticospinal tract | 218 | 18 |
| 15 | Cuneate nucleus | 166 | 168 |
| 16 | Facial nerve | 19 | 219 |
| 17 | Fasciculus retroflexus | 25 | 125 |
| 18 | Fimbria | 211 | 11 |
| 19 | Fornix | 122 | 22 |
| 20 | Fourth ventricle | 118 | 118 |
| 21 | Fundus of striatum | 54 | 154 |
| 22 | Dorsal pallidum (globus pallidus) | 44 | 144 |
| 23 | Habenular commissure | 99 | 199 |
| 24 | Hypothalamus | 250 | 150 |
| 25 | Inferior olivary complex | 113 | 2013 |
| 26 | Internal capsule | 112 | 12 |
| 27 | Interpeduncular nucleus | 157 | 157 |
| 28 | Lateral olfactory tract | 101 | 102 |
| 29 | Lateral septum | 207 | 207 |
| 30 | Lateral ventricle | 57 | 77 |
| 31 | mammillary bodies | 161 | 61 |
| 32 | mammilothalamic tract | 210 | 212 |
| 33 | Medial Lemniscus | 20 | 120 |
| 34 | Medial septum | 53 | 153 |
| 35 | Medulla | 174 | 174 |
| 36 | Midbrain | 194 | 194 |
| 37 | Nucleus Accumbens | 55 | 155 |
| 38 | Olfactory peduncle | 5 | 105 |
| 39 | Olfactory tubercle | 95 | 145 |
| 40 | Optic tract | 216 | 116 |
| 41 | Periaqueductal grey | 10 | 10 |
| 42 | Pons | 187 | 187 |
| 43 | Pontine nucleus | 85 | 185 |
| 44 | Posterior commissure | 100 | 100 |
| 45 | Subiculum | 133 | 131 |
| 46 | Stria medullaris | 225 | 205 |
| 47 | Stria terminalis | 59 | 159 |
| 48 | Striatum | 7 | 17 |
| 49 | Subpendymale zone | 240 | 140 |
| 50 | Superior olivary complex | 124 | 214 |
| 51 | Thalamus | 204 | 4 |
| 52 | Third ventricle | 146 | 146 |
| 53 | Ventral tegmental decussation | 156 | 156 |
| 54 | Cerebellar vermis lobules 1-2 Lingula and ventral central | 32 | 32 |
| 55 | Cerebellar vermis lobule 3: Dorsal central | 233 | 233 |
| 56 | Cerebellar vermis lobules 4-5: culmen | 34 | 34 |
| 57 | Cerebellar vermis lobule 6: declive | 36 | 36 |
| 58 | Cerebellar vermis lobule 7: tuber/folium | 237 | 237 |
| 59 | Cerebellar vermis lobule 8: pyramus | 38 | 38 |
| 60 | Cerebellar vermis lobule 9: uvula | 239 | 239 |
| 61 | Cerebellar vermis lobule 10: nodulus | 40 | 40 |
| 62 | Cerebellar paravermis lobules 4-5: anterior lobule | 90 | 148 |
| 63 | Cerebellar hemisphere lobule 6: simple lobule | 191 | 91 |
| 64 | Cerebellar hemisphere lobule 6: ansiform lobule (crus 1) | 92 | 192 |
| 65 | Cerebellar hemisphere lobule 7: ansiform lobule (crus 2) | 193 | 93 |
| 66 | Cerebellar hemisphere lobule 7: paramedian lobule | 94 | 200 |
| 67 | Cerebellar hemisphere lobule 8: copula pyramidis | 196 | 96 |
| 68 | Flocculus | 97 | 197 |
| 69 | Paraflocculus | 198 | 98 |
| 70 | Trunk of arbor vita | 47 | 47 |
| 71 | Cerebellar vermis WM: lobules 1-2 | 232 | 232 |
| 72 | Cerebellar vermis WM: lobule 3 | 33 | 33 |
| 73 | Cerebellar vermis WM: trunk of lobules 1-3 | 253 | 253 |
| 74 | Cerebellar vermis WM: lobules 4-5 | 234 | 234 |
| 75 | Cerebellar vermis WM: lobules 6-7 | 236 | 236 |
| 76 | Cerebellar vermis WM: lobule 8 | 238 | 238 |
| 77 | Cerebellar vermis WM: trunk of lobules 6-8 | 254 | 254 |
| 78 | Cerebellar vermis WM: lobule 9 | 139 | 139 |
| 79 | Cerebellar vermis WM: lobule 10 | 252 | 252 |
| 80 | Cerebellar paravermis WM: anterior lobule | 21 | 31 |
| 81 | Cerebellar WM: simple lobule | 241 | 251 |
| 82 | Cerebellar WM: crus 1 | 220 | 170 |
| 83 | Cerebellar WM: trunk of simple and crus 1 | 226 | 246 |
| 84 | Cerebellar WM: crus 2 | 229 | 249 |
| 85 | Cerebellar WM: paramedian lobule | 228 | 248 |
| 86 | Cerebellar WM: trunk of crus 2 and paramedian | 175 | 195 |
| 87 | Cerebellar WM: copula | 224 | 244 |
| 88 | Paraflocculus WM | 183 | 243 |
| 89 | Flocculus WM | 167 | 177 |
| 90 | Dentate nucleus | 1 | 201 |
| 91 | Nucleus interpositus | 203 | 3 |
| 92 | Fastigial nucleus | 15 | 206 |
| 93 | Cingulate cortex: area 24a | 24 | 169 |
| 94 | Cingulate cortex: area 24a' | 26 | 171 |
| 95 | Cingulate cortex: area 24b | 27 | 172 |
| 96 | Cingulate cortex: area 24b' | 28 | 173 |
| 97 | Cingulate cortex: area 25 | 29 | 178 |
| 98 | Cingulate cortex: area 29a | 30 | 179 |
| 99 | Cingulate cortex: area 29b | 35 | 182 |
| 100 | Cingulate cortex: area 29c | 37 | 184 |
| 101 | Cingulate cortex: area 30 | 39 | 186 |
| 102 | Cingulate cortex: area 32 | 41 | 188 |
| 103 | Amygdalopiriform transition area | 42 | 189 |
| 104 | Primary auditory cortex | 46 | 208 |
| 105 | Dorsal Secondary auditory cortex | 48 | 217 |
| 106 | Ventral Secondary auditory cortex | 49 | 221 |
| 107 | Caudomedial entorhinal cortex | 50 | 227 |
| 108 | Cingulum | 56 | 231 |
| 109 | Claustrum | 58 | 235 |
| 110 | Piriform area | 60 | 255 |
| 111 | Dorsal Claustrum | 62 | 256 |
| 112 | Dorsal Endopiriform nucleus | 65 | 257 |
| 113 | Dorsal intermediate entorhinal cortex | 67 | 258 |
| 114 | Dorsolateral entorhinal cortex | 69 | 259 |
| 115 | Dorsolateral orbital cortex | 70 | 260 |
| 116 | Dorsal tenia tecta | 71 | 261 |
| 117 | Ectorhinal cortex | 72 | 262 |
| 118 | Frontal cortex: area 3 | 73 | 263 |
| 119 | Frontal association cortex | 74 | 264 |
| 120 | Intermediate nucleus of endopiriform claustrum | 75 | 265 |
| 121 | Insular region: not subdivided | 78 | 266 |
| 122 | Lateral orbital cortex | 79 | 267 |
| 123 | Lateral parietal association cortex | 80 | 268 |
| 124 | Primary motor cortex | 81 | 269 |
| 125 | Secondary motor cortex | 82 | 270 |
| 126 | Medial entorhinal cortex | 83 | 271 |
| 127 | Medial orbital cortex | 84 | 272 |
| 128 | Medial parietal association cortex | 86 | 273 |
| 129 | Piriform cortex | 87 | 274 |
| 130 | Posterolateral cortical amygdaloid area | 88 | 275 |
| 131 | Posteromedial cortical amygdaloid area | 89 | 176 |
| 132 | Perirhinal cortex | 104 | 277 |
| 133 | arietal cortex: posterior area: rostral part | 108 | 278 |
| 134 | Rostral amygdalopiriform area | 110 | 279 |
| 135 | Primary somatosensory cortex | 111 | 280 |
| 136 | Primary somatosensory cortex: barrel field | 117 | 281 |
| 137 | Primary somatosensory cortex: dysgranular zone | 121 | 282 |
| 138 | Primary somatosensory cortex: forelimb region | 126 | 283 |
| 139 | Primary somatosensory cortex: hindlimb region | 127 | 284 |
| 140 | Primary somatosensory cortex: jaw region | 128 | 285 |
| 141 | Primary somatosensory cortex: shoulder region | 129 | 286 |
| 142 | Primary somatosensory cortex: trunk region | 132 | 287 |
| 143 | Primary somatosensory cortex: upper lip region | 134 | 288 |
| 144 | Secondary somatosensory cortex | 135 | 289 |
| 145 | Temporal association area | 136 | 290 |
| 146 | Primary visual cortex | 137 | 291 |
| 147 | Primary visual cortex: binocular area | 138 | 292 |
| 148 | Primary visual cortex: monocular area | 141 | 293 |
| 149 | Secondary visual cortex: lateral area | 142 | 294 |
| 150 | Secondary visual cortex: mediolateral area | 147 | 295 |
| 151 | Secondary visual cortex: mediomedial area | 149 | 296 |
| 152 | Ventral Claustrum | 158 | 297 |
| 153 | Ventral nucleus of the endopiriform claustrum | 160 | 298 |
| 154 | Ventral intermediate entorhinal cortex | 162 | 299 |
| 155 | Ventral orbital cortex | 163 | 300 |
| 156 | Ventral tenia tecta | 165 | 301 |
| 157 | Hippocampal region: CA10r | 336 | 305 |
| 158 | Hippocampal region: LMol | 306 | 307 |
| 159 | Hippocampal region: CA1Rad | 308 | 309 |
| 160 | Hippocampal region: CA2Py | 310 | 311 |
| 161 | Hippocampal region: CA20r | 312 | 313 |
| 162 | Hippocampal region: CA2Rad | 314 | 315 |
| 163 | Hippocampal region: CA3Py Inner | 316 | 317 |
| 164 | Hippocampal region: CA3Py Outer | 318 | 319 |
| 165 | Hippocampal region: CA30r | 320 | 321 |
| 166 | Hippocampal region: CA3Rad | 322 | 323 |
| 167 | Hippocampal region: SLu | 324 | 325 |
| 168 | Hippocampal region: MoDG | 326 | 327 |
| 169 | Hippocampal region: GrDG | 328 | 329 |
| 170 | Hippocampal region: PoDG | 330 | 331 |
| 171 | Hippocampal region: CA1Py | 334 | 335 |
| 172 | Olfactory bulb: glomerular layer | 337 | 345 |
| 173 | Olfactory bulb: external plexiform layer | 338 | 346 |
| 174 | Olfactory bulb: mitral cell layer | 339 | 347 |
| 175 | Olfactory bulb: internal plexiform layer | 340 | 348 |
| 176 | Olfactory bulb: granule cell layer | 341 | 349 |
| 177 | Accessory olfactory bulb: glomerular, external plexiform and mitral cell layer | 342 | 350 |
| 178 | Accessory olfactory bulb: granule cell layer | 343 | 351 |
| 179 | Anterior olfactory nucleus | 344 | 352 |
| 180 | subiculum | 332 | 333 |
| 181 | Medial amygdala | 353 | 355 |
| 182 | Medial preoptic nucleus | 354 | 356 |
